# *Fgf8* dosage regulates jaw shape and symmetry through pharyngeal-cardiac tissue relationships

**DOI:** 10.1101/2022.03.17.484804

**Authors:** Nathaniel Zbasnik, Katie Dolan, Stephanie A. Buczkowski, Rebecca Green, Benedikt Hallgrimsson, Ralph S. Marcucio, Anne M. Moon, Jennifer L. Fish

## Abstract

**Background:** Asymmetries in craniofacial anomalies are commonly observed. With respect to the facial skeleton, the left side is more commonly and/or severely affected than the right. Such asymmetries complicate treatment options. Mechanisms underlying variation in disease severity between individuals as well as within individuals (asymmetries) are still relatively unknown.

**Results:** Developmental reductions in Fibroblast growth factor 8 (Fgf8) have a dosage dependent effect on jaw size, shape, and symmetry. Further, *Fgf8* mutants have directionally asymmetric jaws with the left side being more affected than the right. Defects in lower jaw development begin with an early disruption to Meckel’s cartilage, which is discontinuous and appears as two separate condensations in *Fgf8* mutants. All skeletal elements associated with the proximal condensation are dysmorphic in the mutants, which is exemplified by a malformed and mis-oriented malleus. At later stages, *Fgf8* mutants exhibit syngnathia, which falls into 2 broad categories: bony fusion of the maxillary and mandibular alveolar ridges and zygomatico-mandibular fusion. All of these morphological defects exhibit both inter- and intra-individual variation.

**Conclusions:** We hypothesize that these asymmetries are linked to asymmetries in heart development resulting in higher levels of *Fgf8* on the right side of the face during development, which may buffer the right side to mild developmental perturbations. This mutant mouse is a good model for future investigations of mechanisms underlying human syngnathia and facial asymmetry.

## 1 INTRODUCTION

Among the most common and debilitating human congenital birth defects are those that affect the craniofacial complex [1]. Treatment of craniofacial malformations is often complicated by variation in disease severity [2, 3]. Variation in severity can also occur within individuals, exhibited as left-right asymmetry. Facial asymmetry is typical in healthy populations (nonpathologic asymmetry), with mandibular asymmetries being the most common [4]. In fact, recent studies indicate that more than half of the general population exhibits some degree of mandibular asymmetry and that right-side dominance appears in up to 80% of individuals [4, 5]. Congenital malformations exhibiting asymmetry include facial clefts, hemifacial microsomia, and craniosynostosis. Notably, unilateral clefts of the lip occur on the left side twice as often as clefts on the right side [6]. It is unclear what mechanisms underlie this directional asymmetry.

In a previous study, we found that phenotypic variation in craniofacial disease outcomes can be explained by a non-linear relationship between gene expression and facial shape [7]. The non-linear model posits that reduction in a particular developmental factor (e.g., gene expression) can be buffered until a point, but when molecular levels drop below the buffered level, high levels of phenotypic variance are observed. In other words, when molecular levels are high, differences in expression of 5-10% (or even 50%) do not generate different phenotypes. However, when molecular levels are low, differences of 5-10% cause quantitative differences in morphology (see Fig. 7E). This non-linear model helps to explain why disease phenotypes exhibit differences in severity among affected individuals [7, 8]. A further prediction of this model is that facial asymmetry derives from left-right differences in levels of important developmental factors.

The jaw develops from the growth and fusion of a series of facial primordia composed of neural crest and mesodermal mesenchyme surrounded by several epithelia. The lower jaw forms from the paired mandibular processes of the first pharyngeal arch (PA1), while the upper jaw derives predominantly from the paired maxillary prominences with minor contributions from the frontonasal processes. The lower jaw, or mandible, is subsequently formed by the fusion of the left and right dentary bones which develop largely independently from each other. The relative morphological simplicity of the lower jaw makes it a great system to study phenotypic variation. Additionally, the developmental independence of the left and right halves of the lower jaw allows the effect of subtle differences in developmental perturbations on morphology to be observed.

*Fgf8* is expressed in several tissues important for craniofacial development, including the oral ectoderm of PA1, the lateral ectoderm of the pharyngeal clefts, the foregut endoderm of the pharyngeal pouches, and the anterior heart field mesoderm. To further investigate how *Fgf8* dosage contributes to variation in craniofacial phenotypes, we utilized an allelic series of mutant mice that generates embryos expressing different levels of *Fgf8* during development, including a mild and severe mutant [9]. Quantification of *Fgf8* mRNA levels in E10.5 heads indicates that mice heterozygous for the Neo allele (*Fgf8*^*Neo/+*^) express 90% of WT levels, mice heterozygous for the Delta allele (*Fgf8*^*Δ/+*^) express 60% of WT levels, mice homozygous for the Neo allele (*Fgf8*^*Neo/Neo*^) express 35% of WT levels, and compound mutants (*Fgf8*^*Δ/Neo*^) express 20% of WT levels [7]. Results reported here were generated using these mice, as well as an additional null allele, in which the *Fgf8* coding sequence has been replaced by a LacZ cassette. The two null alleles are functionally equivalent [10] and both are referred to here as “Delta” alleles.

Our previous work quantitatively related differences in *Fgf8* gene expression (molecular variation) to facial shape (morphological variation). Here, we further detail how *Fgf8* dosage contributes to variation in disease severity, focusing on development of the lower jaw, particularly Meckel’s cartilage and its derivatives. We describe not only inter-individual variation in lower jaw defects, but also intra-individual variation. Specifically, *Fgf8* mutants exhibit directional asymmetry with the left side of the jaw more affected than the right. We hypothesize that cardio-pharyngeal developmental interactions contribute to increased *Fgf8* expression on the right side of PA1 during early development. This difference is likely to be small, but significant enough to buffer reductions in *Fgf8* when levels are low (on the high slope of the non-linear curve).

## 2 RESULTS

### 2.1 Reduction in *Fgf8* causes dysmorphic and asymmetric jaws

To investigate the role of *Fgf8* dosage on jaw size, we first evaluated the lower jaws of neonatal (P0) mice from the allelic series. Dentary bones in mice heterozygous for *Fgf8* are phenotypically normal, suggesting that significant reductions in *Fgf8* dosage are buffered in jaw development. However, in both mutant genoytpes, the dentary bones are hypoplastic and dysmorphic to varying degrees (Fig. 1; SupFig1). Morphological defects are concentrated proximally. The distal most portion of the dentary is unaffected, with rostral process and incisors intact, consistent with the distal region of the jaw being *Fgf8* independent [11]. Mandibles from *Fgf8*^*Δ/Neo*^ neonates exhibit severe phenotypes, while neonatal *Fgf8*^*Neo/Neo*^ mandibles are comparatively mild. In the mildest *Fgf8*^*Neo/Neo*^ mutants, only the coronoid process is missing while the rest of the dentary bone appears normal. However, in some *Fgf8*^*Neo/Neo*^ individuals, more severe phenotypes are apparent, including fusion of the dentary to the maxilla (see SupFig1 for range of variation). When syngnathia occurs in *Fgf8*^*Neo/Neo*^ jaws, it is typically unilateral (11/34 *Fgf8*^*Neo/Neo*^ neonates exhibit unilateral fusion on the left side; 1/34 had bilaterally fused jaws). In *Fgf8*^*Δ/Neo*^ neonates, syngnathia is always bilateral (20/20).

**Figure 1:**
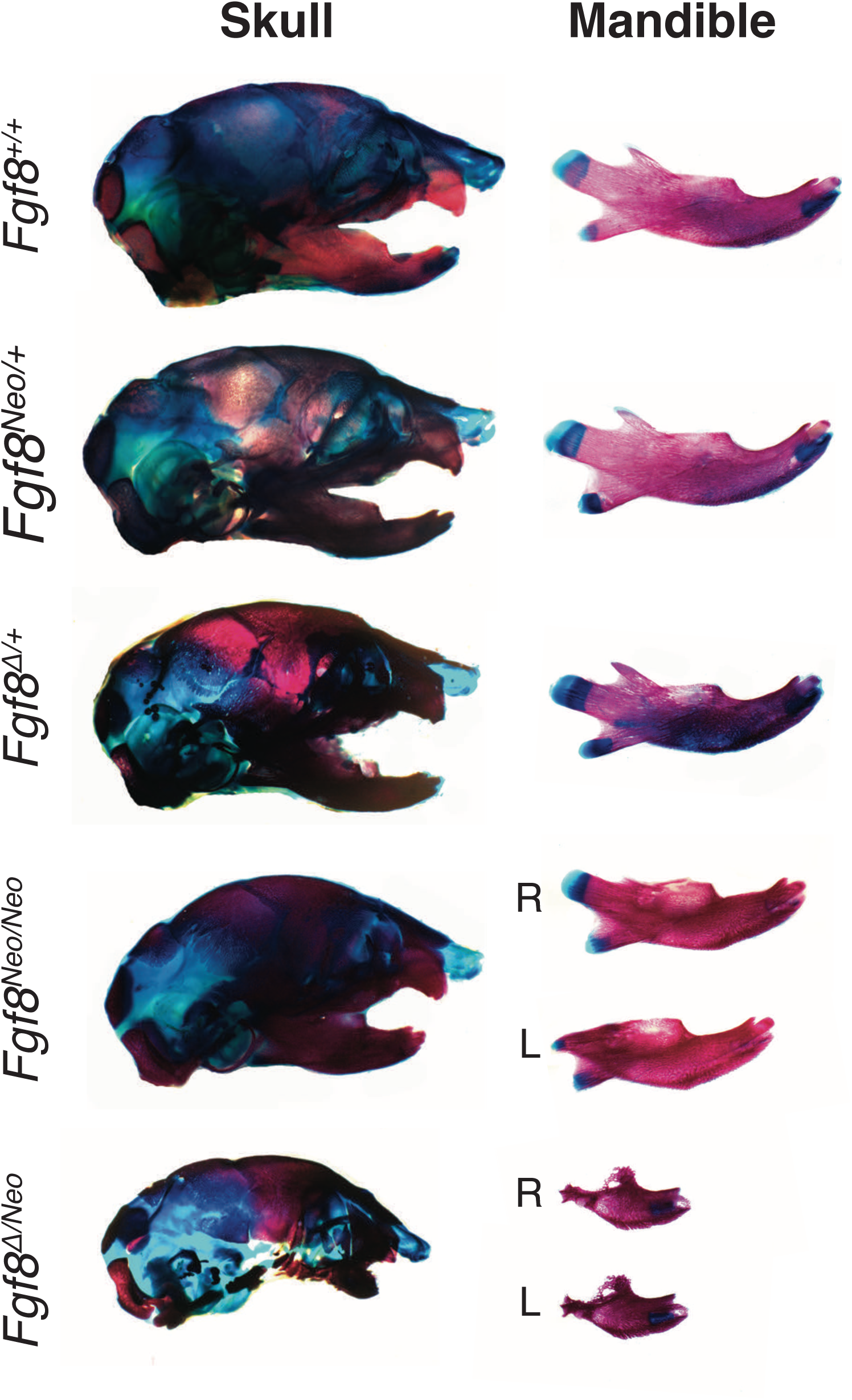
Mandibles exhibit dosage sensitivity and directional asymmetry. Neonatal skulls (left) and isolated dentary bones (right) differentially stained for bone (red) and cartilage (blue). Representative images are shown for *Fgf8*^*+/+*^ (WT), *Fgf8*^*Neo/+*^, *Fgf8*^*Δ/+*^, *Fgf8*^*Neo/Neo*^, and *Fgf8*^*Δ/Neo*^ neonates.

We observed two broad categories of syngnathia: 1) fusion of the maxillary zygomatic process and/or jugal to the dentary and 2) bony fusion of the maxilla and dentary (zygomatic process and jugal bones may be absent) (Fig. 2). These two categories were most obvious in the *Fgf8*^*Δ/Neo*^ mandibles, where both types of fusion occur (Fig. 2G-O). In *Fgf8*^*Neo/Neo*^ neonates, we only observed fusion of upper and lower jaw elements on the dorso-lateral side of the dentary, posterior to the molar alveolus. This fusion sometimes appears in a mild form with a jugal-zygomatic process articulation apparent (black asterisks in SupFig1). In other cases, the fusion is complete and forms a wide bony connection (green asterisks in SupFig1). In *Fgf8*^*Δ/Neo*^ neonates, fusion of the zygomatic process occurs more distal and lateral on the dentary than is observed in *Fgf8*^*Neo/Neo*^ neonates. Further, *Fgf8*^*Δ/Neo*^ neonates typically exhibit bony fusion between the maxilla and dentary that connects the entire dorsal side of the molar alveolar region of the dentary to the upper jaw (Fig. 2J-O). Notably, even in the most severe cases of fusion, rudimentary condylar and angular processes are typically present albeit dysmorphic (Fig. 2; SupFig1).

**Figure 2:**
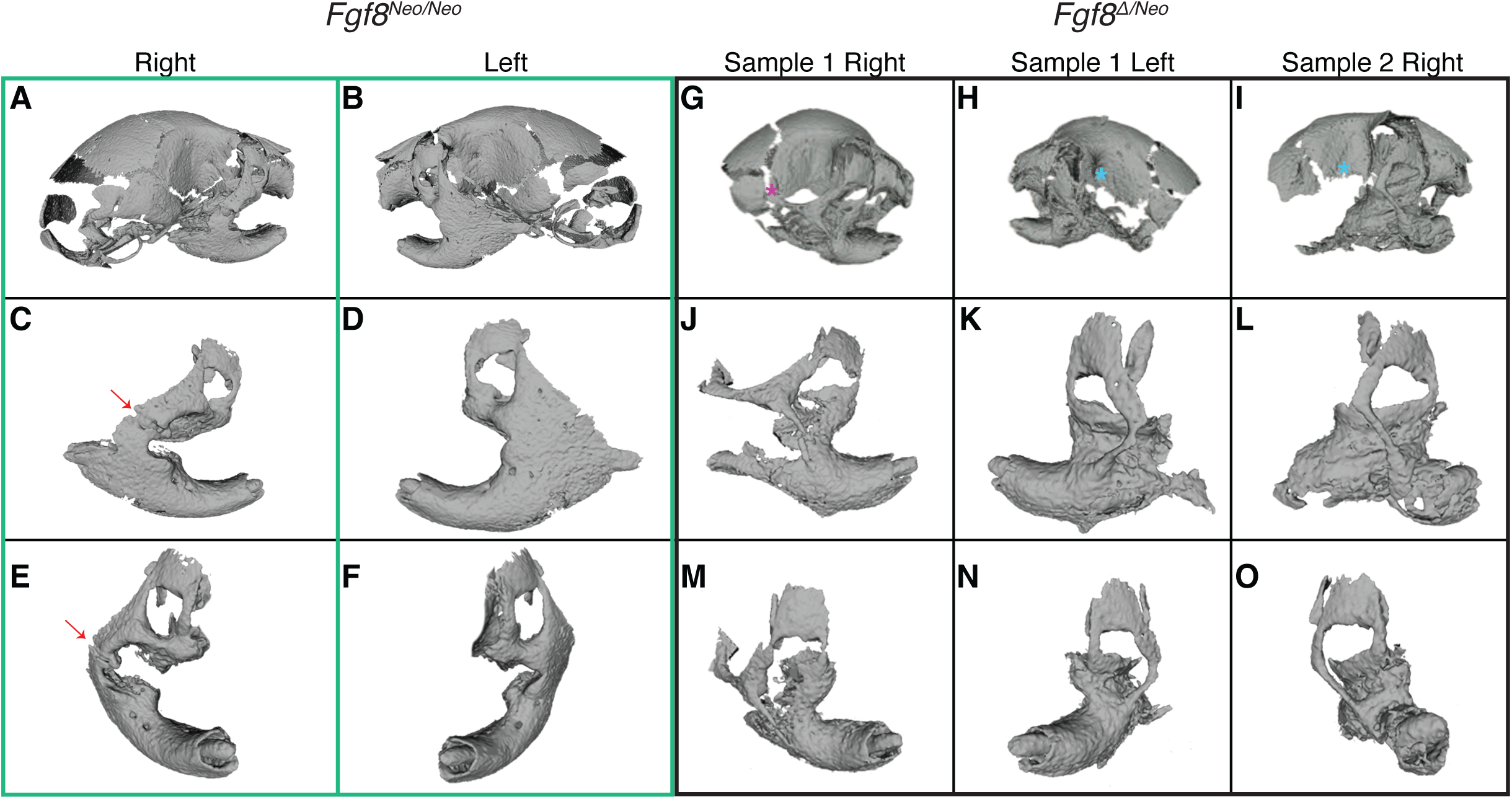
*Fgf8*-mediated jaw fusion is variable. Three-dimensional reconstructions from microCTs of A-F) *Fgf8*^*Neo/Neo*^ and G-O) *Fgf8*^*Δ/Neo*^ neonatal skulls and jaws. Fusion of *Fgf8*^*Neo/Neo*^ jaws occurs between a broadened zygomatic arch at the lateral side of the dentary, however in *Fgf8*^*Δ/Neo*^ jaws an additional fusion is present between the dentary and maxillary process. The zygomatic fusion in *Fgf8*^*Δ/Neo*^ jaws occurs as a thinner bony element that is more distal and lateral on the dentary than what is observed in *Fgf8*^*Neo/Neo*^ jaws. Panels C-F and J-O show individual outer, then frontal, aspects of a single dentary of the corresponding skull (shown in the upper panels). A-F) The zygomatic fusion is more severe on the left side than the right side in *Fgf8*^*Neo/Neo*^ (arrows in C and E point to unfused zygomatic contact). Although the overall fusion is similar on both sides of the jaw in *Fgf8*^*Δ/Neo*^, the squamosal process can be G) present (pink asterisk) or H-I) absent (turquoise askterisks). Data shown are representative of 8 *Fgf8*^*Neo/Neo*^ and 7 *Fgf8*^*Δ/Neo*^ neonatal skulls.

Despite the wide variation in severity of dysmorphology between individuals, we also noted that within individuals, the left dentary bone was typically more severely disrupted than the right, which was especially apparent in *Fgf8*^*Neo/Neo*^ individuals. To further assess directional asymmetry, we measured the length of isolated neonatal dentary bones from WT and *Fgf8*^*Neo/Neo*^ individuals, comparing right versus left sides (SupFig2). Our measurements indicate that the dentary bones of *Fgf8*^*Neo/Neo*^ neonates are shorter than in WT or *Fgf8*^*Neo/+*^ individuals, and that the left side is on average shorter than the right. In contrast, skeletal elements from the limbs, while smaller in *Fgf8*^*Neo/Neo*^ mutants, are symmetrical (SupFig2). Taken together, these data indicate that below an initial buffered level, *Fgf8* has a dosage effect on jaw size and symmetry.

### 2.2 Loss of coronoid is independent of temporalis

To further assess proximal defects of the mandible, we evaluated microCTs of neonatal skulls (Fig. 3). In all *Fgf8*^*Neo/Neo*^ mutants, the coronoid is absent or severely reduced bilaterally (SupFig1; Fig. 3C). Notably, the temporalis muscle attaches to the dorsal region of the dentary where a rudimentary coronoid is sometimes present (Fig. 3B). Since it has been shown that mechanical load from the temporalis muscle maintains growth and regulates size of the coronoid process [12, 13], we calculated the volume of the temporalis muscle from 3D reconstructions of microCT scans of neonatal mouse heads to determine a potential role for the temporalis muscle in coronoid asymmetry. We found that the muscle volume is smaller in mutants than in WT, however, there is no bilateral difference in size (Fig. 3D; n=6 for WT, n=3 for mutant). These data suggest that mechanical signals from the temporalis are not responsible for morphological asymmetry and are unlikely to explain the absence of the coronoid.

**Figure 3:**
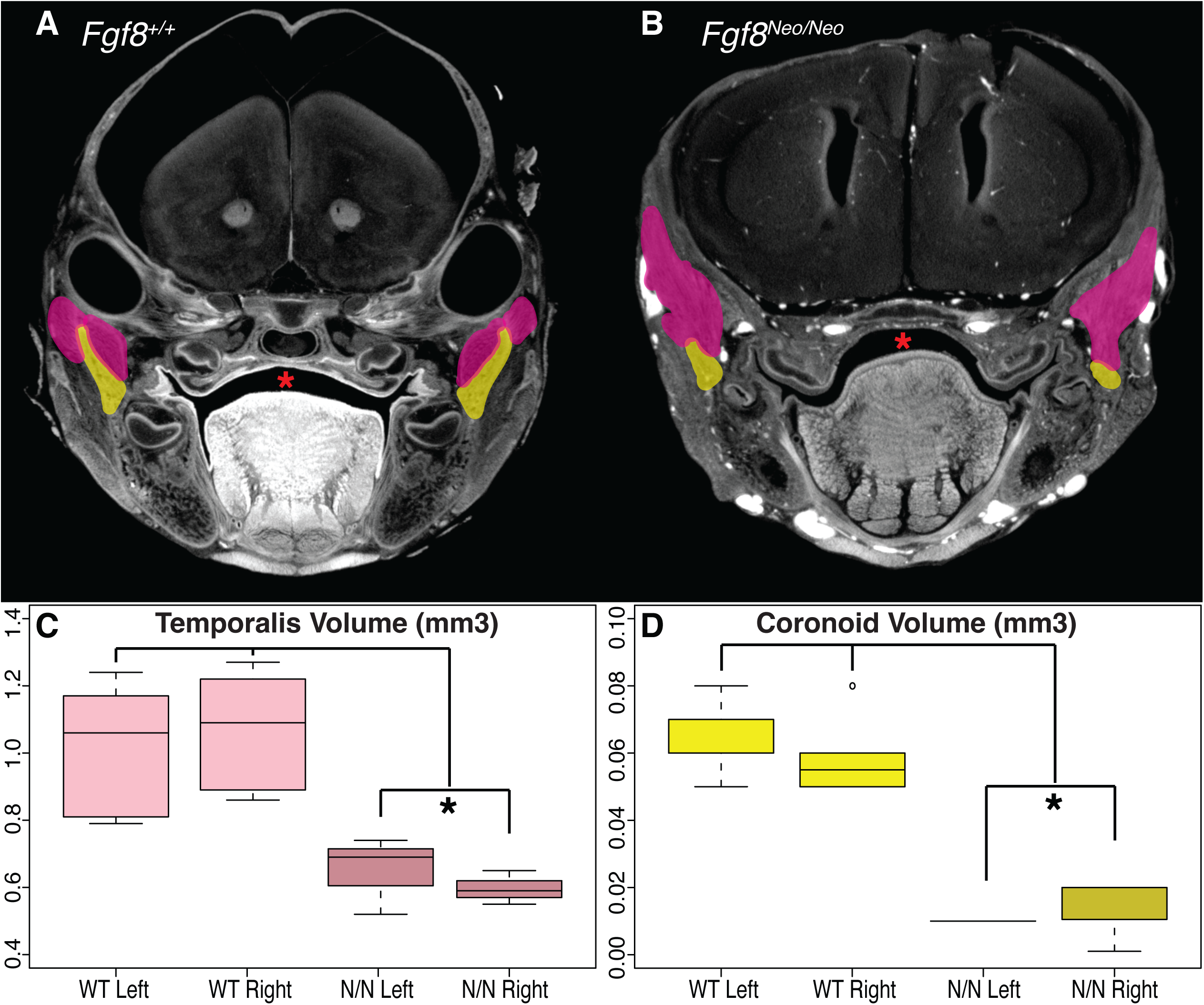
*Fgf8*^*Neo/Neo*^ mutants have small temporalis muscles and absent coronoid processes. Cross-sections from micro-CT scans of A) *Fgf8*^*+/+*^ (WT) and B) *Fgf8*^*Neo/Neo*^ neonatal skulls highlighting the coronoid (yellow) and temporalis muscles (pink). C) Coronoid volume and D) temporalis volume are represented in mm^3^. The section where the coronoid was the largest was chosen for each sample in A and B. The temporalis appears larger in this particular section of the *Fgf8*^*Neo/Neo*^ embryo because shape changes in the muscle related to the smaller jaw and proximal shift in the location of the coronoid relative to the rest of the skull condense the temporalis. Red asterisks highlight normal palate formation in WT and the cleft palate in *Fgf8*^*Neo/Neo*^. *P-value < 0.05, Tukey’s HSD Test. n=6 for WT, n=3 for *Fgf8*^*Neo/Neo*^.

### 2.3 *Fgf8* mutants exhibit discontinuous Meckel’s cartilage

In addition to hypoplasia of the jaw bones, disruptions to Meckel’s cartilage (MC) exhibit *Fgf8* dosage sensitivity. In both *Fgf8*^*Neo/Neo*^ and *Fgf8*^*Δ/Neo*^ mutants, MC is discontinuous near its proximal end where it appears as two separate condensations forming proximal and distal segments (Fig. 4 and SupFig 3). This disruption to MC is observed in 100% of mutant embryos of both genotypes (observed in 6/6 embryos of each genotype) and occurs early in skeletal development, as it is apparent by E14.5. In the mildest mutants, this discontinuity is unilateral, occurring only on the left side (observed in 1 embryo), while in individuals where the discontinuity is bilateral, the gap is typically greater on the left side (data not shown and SupFig3). These data are consistent with our observation of directional asymmetry in the dentary bones. Further, the distance between the distal and proximal segments of MC is greater in *Fgf8*^*Δ/Neo*^ than it is in *Fgf8*^*Neo/Neo*^ embryos, suggesting an *Fgf8* dosage effect (SupFig3I,L).

**Figure 4:**
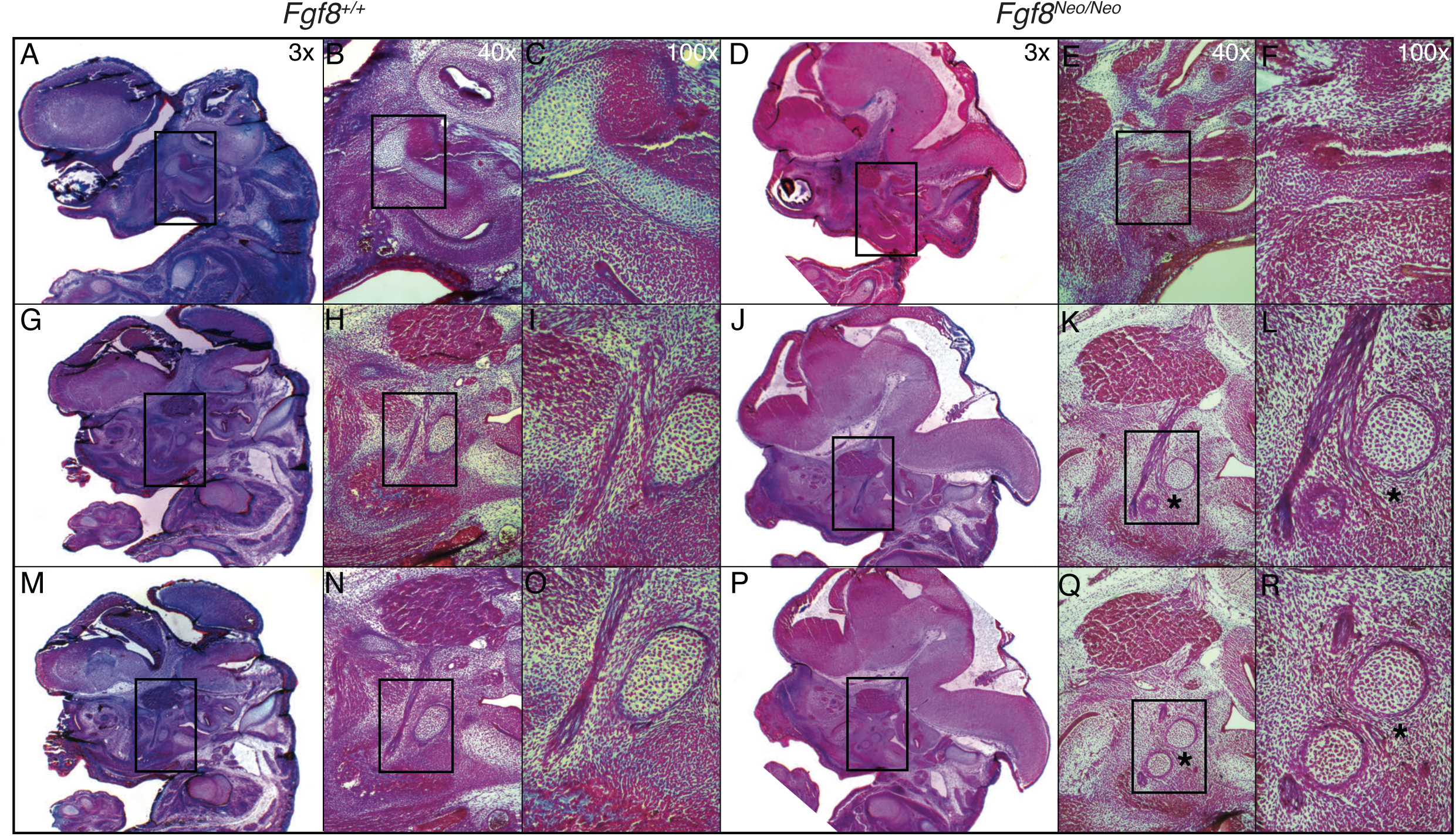
Meckel’s cartilage of *Fgf8*^*Neo/Neo*^ mutants is separated into two non-continuous condensations. Trichrome stained cross-sections of *Fgf8*^*+/+*^ (WT) and *Fgf8*^*Neo/Neo*^ E14.5 embryonic skulls. Sections are arranged from lateral (top row) to medial (bottom row). Laterally, Meckel’s cartilage (MC) in A-C) *Fgf8*^*+/+*^ and D-F) *Fgf8*^*Neo/Neo*^ is connected to the malleus in the proximal region. G-L) Once the trigeminal branches into the Lingual and Inferior Alveolar Nerve, MC is discontinuous in *Fgf8*^*Neo/Neo*^ creating a proximal and distal segment. M-R) MC is proximal to the branching nerve(s) in *Fgf8*^*+/+*^ while in *Fgf8*^*Neo/Neo*^ the distal segment is between the branches. Asterisks denote the separation of the proximal and distal condensations of MC in *Fgf8*^*Neo/Neo*^. Distal Condensation (DC), Inferior Alveolar Nerve (IA), Malleus (M), Lingual Nerve (LN), Proximal Condensation (PC), Trigeminal Ganglion (TG). Discontinuity at E14.5 was seen bilaterally in 2/2 *Fgf8*^*Neo/Neo*^ heads.

In all cases, the discontinuity in MC consistently occurs at the proximal end of the cartilage. In younger embryos (E14.5), MC is separated at the point where the trigeminal nerve branches into the Lingual and Inferior Alveolar Nerve (Fig. 4K,L). In neonatal (P0) embryos, this discontinuity can be observed behind the developing molars, in the proximal plane of the lower jaw near where the coronoid would normally form. Related to this discontinuity, the most proximal end of MC fails to extend medially and instead curves laterally (SupFig3).

### 2.4 Meckel’s cartilage defects contribute to abnormalities of the middle ear and jaw joint

*Fgf8* reductions result in a truncated and discontinuous Meckel’s cartilage. As a result, there is rostral shift of the jaw joint and middle ear relative to the rostral-caudal axis of the head. The mutant skulls exhibit increased cranial base flexion relative to WT (SupFig3A-C). This flexion results in the cochlea lying behind (caudal and ventral) the middle ear. Further, the middle ear is rostral-caudally compressed such that the malleus is lateral to the incus and stapes at the oval window of the otic vesicle rather than lying distal to the oval window (Fig.5M-P; SupFig4).

**Figure 5:**
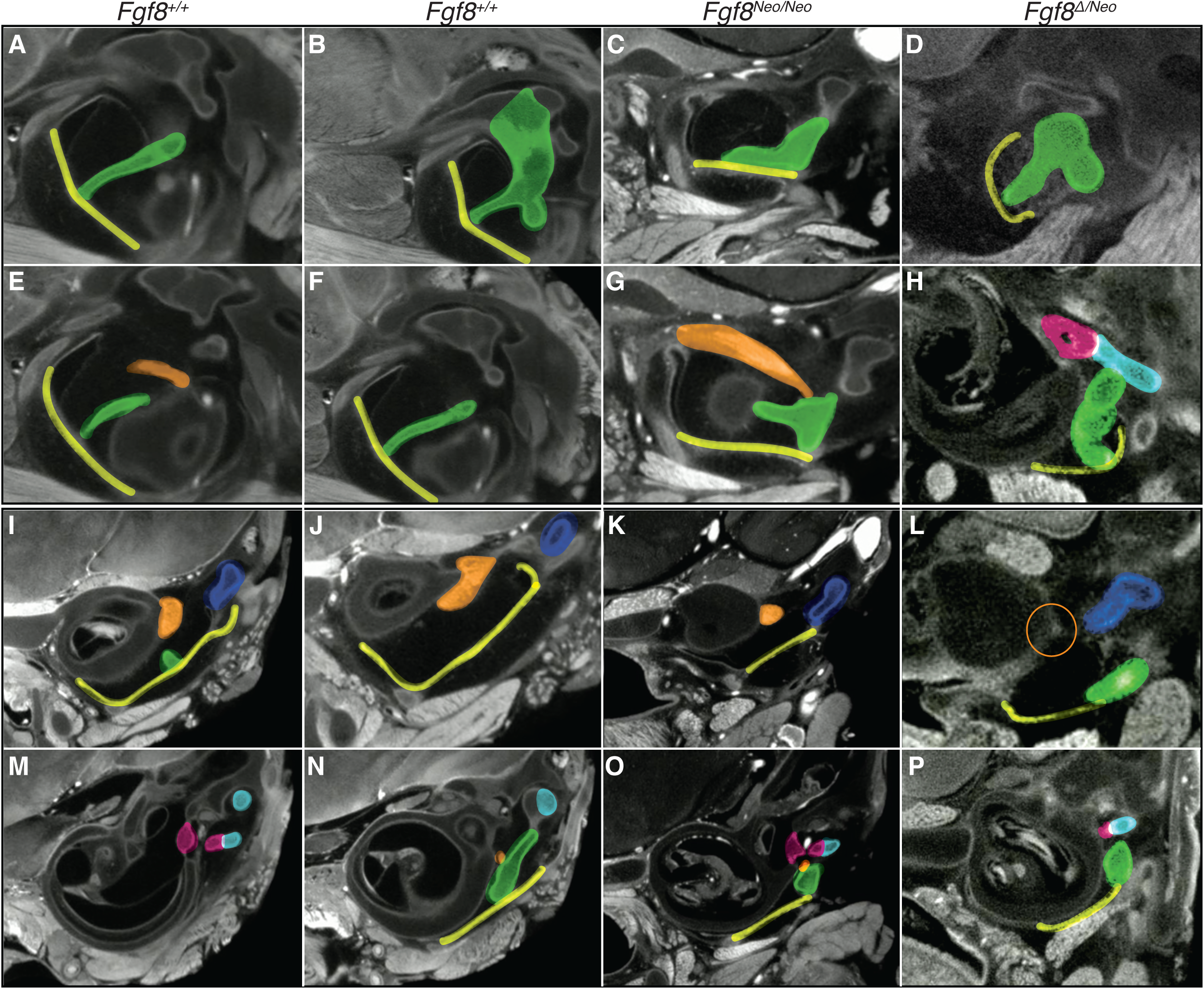
*Fgf8* reductions condense the malleus and middle ear. Micro-CT scans of the middle ear of *Fgf8*^*+/+*^ (WT; 2 left columns), *Fgf8*^*Neo/Neo*^ (3^rd^ column) and *Fgf8*^*Δ/Neo*^ (4^th^ right column) neonatal skulls. Rows 1,2 are oblique sections through the malleus and rows 3,4 are coronal sections through similar regions. WT sections in the first column are proximal to WT sections in column 2. Distal sections are shown in rows 1 and 3; more proximal sections of each sample are shown in rows 2 and 4. B) The tympanic membrane (yellow) creates a V shape with the malleus (green). C,D) The V shape is lost within *Fgf8*^*Neo/Neo*^ and *Fgf8*^*Δ/Neo*^ and the tympanic membrane (TM) is short and linear. E,F) In WT, the manubrium of the malleus is shown where it contacts the TM. In G,H) *Fgf8*^*Neo/Neo*^ the manubrium is dorsally rotated away from the membrane and is altered in shape in *Fgf8*^*Neo/Neo*^ and *Fgf8*^*Δ/Neo*^. I,J) The tympanic cavity is large separating the middle ear and cochlea, but in K) *Fgf8*^*Neo/Neo*^ the cavity is reduced in size and becomes smaller in L) *Fgf8*^*Δ/Neo*^. I,J) Size reductions are similar for the tensor tympani muscle (orange) where the muscle is reduced in K) *Fgf8*^*Neo/Neo*^ and possibly absent in L) *Fgf8*^*Δ/Neo*^ mutants (orange circle). N) This muscle attaches to the malleus at the point where the incus (light blue), Meckel’s cartilage (dark blue), and malleus are together. O) In *Fgf8*^*Neo/Neo*^ the muscle still connects to the malleus but is ventral to the connection of the stapes (pink) and incus. O,P) The malleus appears to dorso-ventrally condense and curve under the stapes and incus. M) Relative to the inner-ear shape in WT, the connection site of the stapes and incus of O) *Fgf8*^*Neo/Neo*^ and P) *Fgf8*^*Δ/Neo*^ is shifted distally above the dorsally condensed malleus. M,O,H) The stapes appears morphologically normal in all dosage levels. External Auditory Canal (EAC), Inner Ear (IE), Manubrium of Malleus (MM), Tympanic Cavity (TC). Representative images are shown for *Fgf8*^*+/+*^ (WT) 2; *Fgf8*^*Neo/+*^ 4 (not shown, but similar to WT); *Fgf8*^*Neo/Neo*^ *3; Fgf8*^*Δ/Neo*^ 1.

In mutant embryos, the malleus is malformed and mis-oriented (Fig. 5; SupFig4). The head of the malleus is reduced in *Fgf8*^*Neo/Neo*^ neonates and in *Fgf8* ^*Δ/Neo*^ mutants it is nearly lost. This contributes to the articulation between malleus and incus being severely reduced in *Fgf8* ^*Δ/Neo*^ mutants (SupFig4). In contrast, the manubrium of the malleus is enlarged and rotated dorsally. As a consequence, the tympanic membrane contacts the malleus at the process brevis in addition to the manubrium. The enlarged process brevis of the malleus lies ventrally on a shortened and straight tympanic membrane, whereas in WT, the manubrium of the malleus connects to the middle of the tympanic membrane at a focal point generating a V-shape in the membrane (Fig. 5A-B, D-F). Additionally, the tensor tympani muscle is smaller in *Fgf8*^*Neo/Neo*^ mutants and absent in *Fgf8* ^*Δ/Neo*^ mutants (Fig. 5H,L).

The compression of the middle ear is associated with a reduction in size of the tympanic cavity in *Fgf8* mutants. In *Fgf8*^*Neo/Neo*^ mutants, this reduced tympanic cavity contributes to the malleus contacting the cochlea. In *Fgf8* ^*Δ/Neo*^ mutants, the tympanic cavity is severely reduced and sometimes absent (SupFig5A-D). When present, the tympanic cavity is separated from the cochlea where the malleus meets the tympanic membrane. In some *Fgf8* ^*Δ/Neo*^ mutants, the tympanic cavity opens into the oral cavity (SupFig5I).

We also observed an *Fgf8* dosage effect on jaw joint formation. In mammals, the jaw articulates at the temporomandibular joint (TMJ) between the condylar process of the dentary and the glenoid fossa of the squamosal bone. In WT (*Fgf8*^*+/+*^) neonatal mice, the TMJ is characterized by an articular disc sitting below the glenoid fossa creating a defined joint cavity (SupFig5E,G). However, in *Fgf8*^*Δ/Neo*^ neonates, the glenoid fossa, condylar process and joint cavity are malformed, and the articular disc is absent (SupFig5F,H).

### 2.5 Apoptosis of neural crest is not asymmetric

Reductions in *Fgf8* have previously been reported to increase apoptosis in migrating and post-migratory neural crest [14–17]. Such reductions in neural crest numbers have been hypothesized to contribute to the hypoplasia associated with craniofacial defects in *Fgf8* mutant mice. We also evaluated apoptosis in E9.5 and E10.5 *Fgf8*^*Neo/Neo*^ heads to investigate if any asymmetry was present that might contribute to the directional asymmetry we observe in the lower jaw. In WT embryos, apoptosis appears as an isolated cluster in the proximal hinge region of PA1, where apoptosis is known to occur in the developing epibranchial placodes [18]. In contrast, in *Fgf8*^*Neo/Neo*^ embryos, apoptosis occurs in large streams of neural crest migrating from the neural tube. However, apoptosis in PA1 of *Fgf8*^*Neo/Neo*^ embryos is minor (SupFig6). Previously, Frank and colleagues (2002) obtained similar results from a different hypomorphic *Fgf8* mouse line [17]. In contrast, in Fgf8;Nestin-Cre embryos, in which *Fgf8* is completely absent in the oral ectoderm, cell death in PA1 is extensive and the mandibular arch is severely hypoplastic by E10.5 [11]. These data suggest that sufficient *Fgf8* levels exist in the oral ectoderm of *Fgf8*^*Neo/Neo*^ embryos to maintain survival of post-migratory neural crest. Notably, we find no evidence suggesting that left-right asymmetry in apoptosis is a driving factor in mandibular asymmetry.

### 2.6 Reduction in *Fgf8* disrupts PA1 patterning

The alterations to the distribution of cell death in E9.5 and E10.5 embryos discussed above suggested that early embryonic patterning may be disrupted when *Fgf8* dosage is reduced. To further assess alterations to PA1 patterning, we investigated the expression of several genes known to be important to early patterning of the jaw. At E10.5, the first pharyngeal arch is typically well developed with a deep invagination of the oral ectoderm separating the maxillary (mx) and mandibular (md) portions of PA1. However, in *Fgf8* ^*Δ/Neo*^ embryos, the oral ectoderm is severely reduced and the hinge region of PA1 is underdeveloped such that the maxillary and mandibular processes are only separated at the most distal end of the arch (Fig. 6A-D). Further, the pharyngeal cleft, which normally separates the first and second arches, fails to extend in *Fgf8* ^*Δ/Neo*^ embryos. As a consequence, the mesenchyme of PA1 and PA2 is continuous (yellow arrows in Fig. 6C,D).

**Figure 6:**
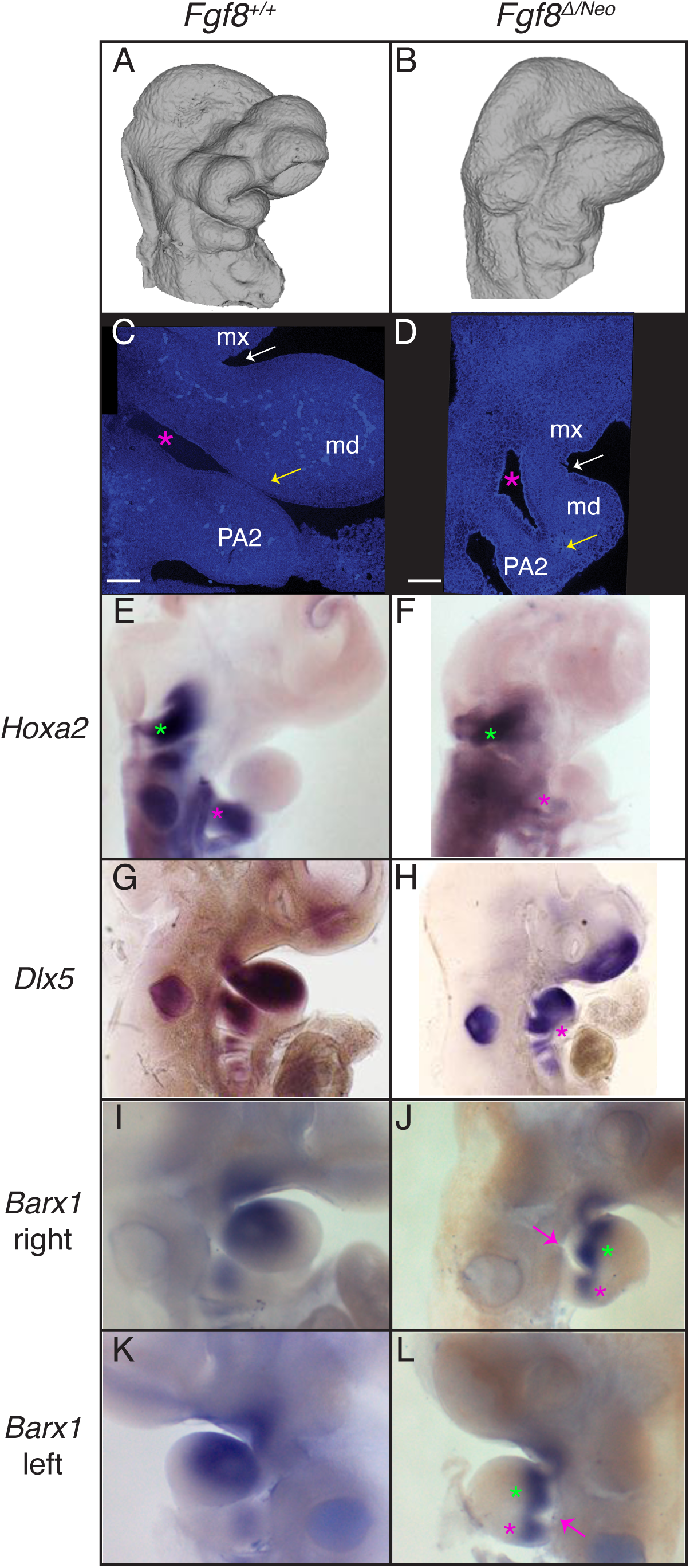
Reductions in *Fgf8* dosage alter pharyngeal tissue relationships and patterning. A,B) Lateral views of microCT images of E10.5 A) *Fgf8*^*+/+*^ (WT) and B) *Fgf8*^*Δ/Neo*^ embryos show reduction of the first pharyngeal arch (PA1) in the mutants. Lateral views of nuclear staining of E10.5 C) *Fgf8*^*+/+*^ (WT) and D) *Fgf8*^*Δ/Neo*^ embryos show alterations to the mandibular (md) and maxillary (mx) portions of PA1. The oral ectoderm does not extend into the hinge region of PA1 (white arrow). The first pharyngeal cleft (green asterisk) fails to extend in the mutants and the first and second arches fail to separate distally (yellow arrow). Whole-mount *in situ* hybridization in E10.5 embryos for (E,F) *Hoxa2*, (G,H) *Dlx5*, and (I-L) *Barx1* show alterations to gene expression in *Fgf8*^*Δ/Neo*^ mutants. In WT, Hoxa2 expression in PA2 (pink asterisk in E) is equivalent to that in the hindbrain (green asterisk), whereas in *Fgf8*^*Δ/Neo*^ embryos, *Hoxa2* is reduced in PA2 (pink asterisk in F). *Dlx5* and *Barx1* expression domains are continuus across PA1 and PA2 where the arches are abnormally continugous (pink asterisks in H,J,L). Additionally, *Barx1* is split into 2 domains in the proximal portion of PA1 (green asterisks). The cleft on the left side of the embryo is more underdeveloped on the left side of the mutant compared to the right (pink arrow). All in situs were performed on at least 2 embryos per genotype.

We found that the expression of *Hoxa2* is severely reduced in PA2 in *Fgf8* ^*Δ/Neo*^ embryos (Fig. 6E,F). This PA1-like expression pattern may be related to its altered association to signaling centers in the pharyngeal arches. Within PA1, we find that *Dlx5* and *Barx1* are also altered in *Fgf8* ^*Δ/Neo*^ E10.5 embryos (Fig. 6G-J). *Dlx5* does not extend past the hinge region in mutant embryos (green arrows in Fig. 6G,H) and is continuous across PA1 and PA2 in the region where the cleft fails to extend (pink asterisk in Fig. 6H). *Barx1* expression is reduced in *Fgf8* ^*Δ/Neo*^ embryos and appears to bifurcate at the distal end of its expression domain (greens asterisk in Fig. 6J,L). Similar to *Dlx5, Barx1* expression is continuous across PA1 and PA2 (pink asterisks in Fig. 6J,L). Further, these embryos show that failure of cleft extension on the left side is more severe than on the right side (pink arrows in Fig. 6J,L). These data are consistent with an alteration to patterning of the pharyngeal arches in *Fgf8* mutants.

### 2.7 *Fgf8* is expressed asymmetrically in the developing heart

*Fgf8* is expressed in several focal areas of the pharyngeal arches, specifically the proximal oral ectoderm overlying, the lateral ectoderm rostral to the pharyngeal clefts, and the endodermal pouches, where it is critical to survival and patterning of neural crest cells [9, 11, 19]. Expression in these domains, especially the oral ectoderm, may exhibit fluctuating asymmetry in early pharyngeal arch development, however, it equilibrates bilaterally by E10.5 [20]. Therefore, we do not consider *Fgf8* expression in these domains to be a causative factor underlying directional asymmetry in the mandible. However, *Fgf8* expression is bilaterally asymmetric in the anterior heart field mesoderm precursors of the outflow tract [21]. This asymmetry is a byproduct of heart looping which shifts the outflow tract to the right side of the pharyngeal region, adjacent to the first and second pharyngeal arches.

To further characterize *Fgf8* expression in this domain relative to the developing first arch, we performed fluorescent *in situ* hybridization for *Fgf8* on E9.5 embryos. We found that ectoderm overlying the developing heart contacts the first arch ectoderm on the right side, but not the left. Instead, on the left side of the embryo, regions of the heart approach the frontonasal process (Fig. 7A-D). *Fgf8* expression within the developing heart at early developmental stages may, therefore, contribute to PA1 development. These tissue relationships provide a possible explanation for directional asymmetry of the jaw, via increased Fgf8 levels on the right side of the embryo at a stage that is critical for mandibular patterning.

**Figure 7:**
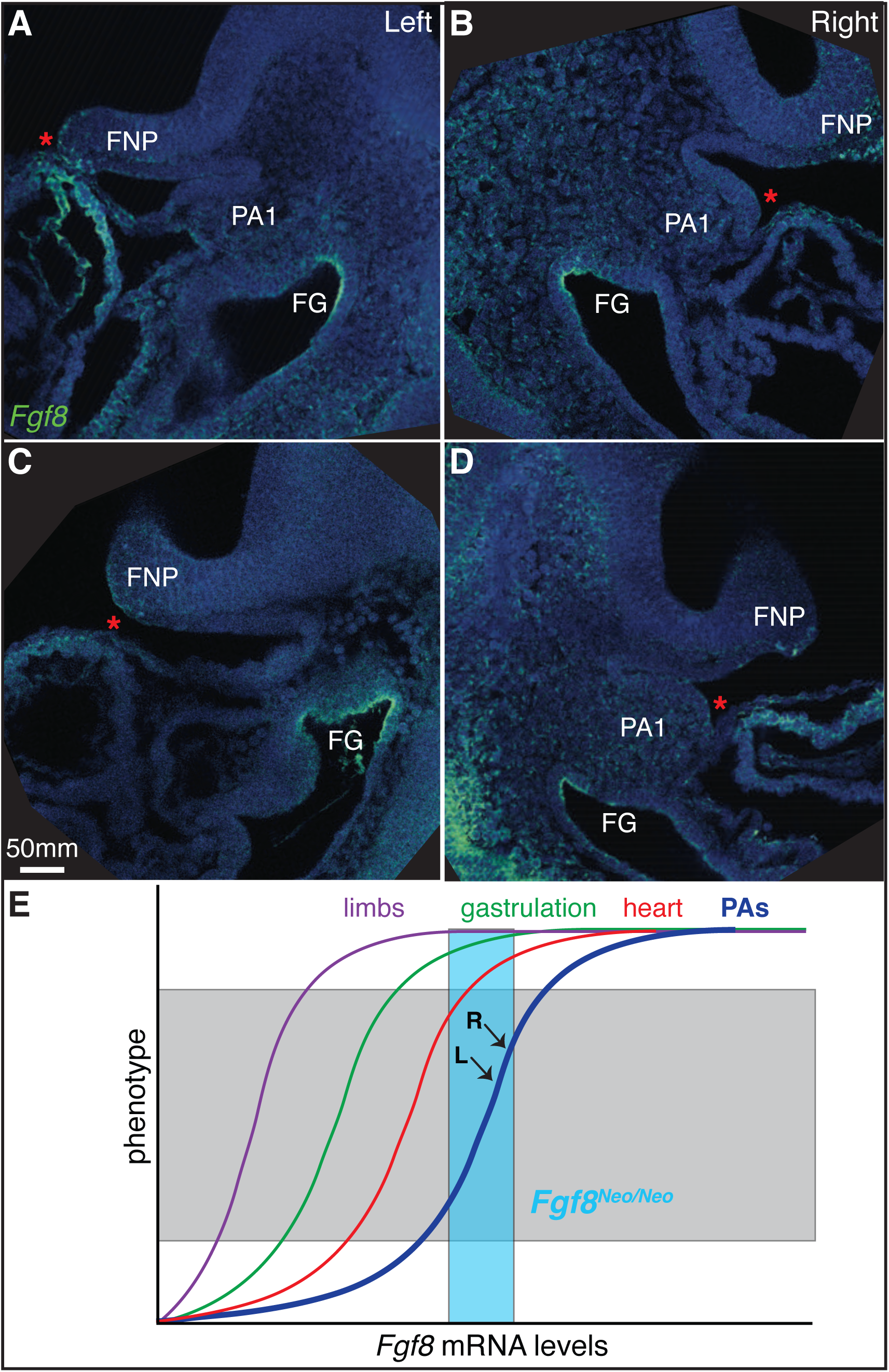
*Fgf8* is expressed asymmetrically in the developing heart. A-D) Saggital sections of fluorescent in situ hybridization of *Fgf8* (green) in E9.5 embryos. Left (A,C) and right (B,D) sides are shown for two separate individuals. Sections shown are those where the developing heart tube comes closest to the first pharyngeal arch (PA1) for each side. Nuclei are counterstained with dapi (blue). E) Model for non-linear relationships for Fgf8 levels and tissue morphogenesis. Teal box represents *Fgf8* levels in *Fgf8*^*Neo/Neo*^ mice. L and R indicate levels in left and right sides of PA1 (see text for more details). Frontonasal process (FNP), foregut (FG).

We did further test this hypothesis by investigating mandibular asymmetry in tissue-specific *Fgf8* knock-outs (SupFig8). Loss of *Fgf8* in the ectoderm using an Ap2α-CRE generates a severely truncated jaw similar to that reported for Fgf8;Nestin-Cre, with some fluctuating asymmetry. Removal of *Fgf8* from the mesoderm using MesP1-CRE had no obvious affect jaw morphology. Because of buffering effects, it’s possible that directional asymmetry only occurs when *Fgf8* is reduced globally. If so, a tissue-specific knock-out in a global hypomorphic background would be required to test this hypothesis. Such a mouse model is not currently available.

## 3 Discussion

### 3.1 Jaw defects are caused by disruptions to pharyngeal patterning

To investigate the impact of *Fgf8* dosage on jaw morphology, we examined the morphology of the lower jaw in mice expressing various levels of *Fgf8* during development. The allelic series of mice we used includes two mutant genotypes, *Fgf8*^*Neo/Neo*^ (mild mutants) and *Fgf8* ^*Δ/Neo*^ (severe mutants). These two genotypes are distinguished by their average level of *Fgf8* expression (roughly 35% vs 20%, respectively) as well as the overall severity of defects. Nevertheless, individuals within each mutant genotype also express variable phenotypes (Figs. 1, 2; SupFig1). For example, the mildest *Fgf8*^*Neo/Neo*^ neonates have relatively normal mandibles that only lack a coronoid process. More severe *Fgf8*^*Neo/Neo*^ neonates exhibit unilateral fusion of the upper and lower jaws. Overall, we find that reductions in *Fgf8* expression below about 40% of WT expression levels result in truncated and dysmorphic lower jaws.

The morphological defects observed in mutant embryos are focal, being localized to the proximal portion of the jaw. Specifically, Meckel’s cartilage is bifurcated near the TMJ, and all derivatives of Meckel’s cartilage proximal to the point of bifurcation are dysmorphic. The distal end of Meckel’s cartilage and its derivatives, including the rostral process and incisor portion of the dentary are grossly morphologically normal, although reduced in size to varying degrees. Proximal defects of Meckel’s cartilage are exemplified by a dysmorphic malleus (Figs. 4, 5; SupFig4). Defects of the middle ear are compounded by increased cranial base flexion which compresses the middle ear on the rostral-caudal axis (SupFig3).

Although the most severe mutant mandibles are small, many mild mutants have relatively normal dentaries in size and shape, lacking only a coronoid process. These mild mutants maintain muscle attachment (Fig. 3), and are therefore not likely to be caused by deficiencies in mechanical signaling [13]. Additionally, despite an increase in apoptosis in migrating neural crest, *Fgf8* mutants nonetheless have well-populated pharyngeal arches. Cell death is observed in migrating neural crest, but is not detected within PA1 at E9.5 or E10.5 (SupFig6). Since neural crest are not regionally specified until E10.5 [19], this suggests that the available neural crest should respond to appropriate patterning signals if present. Further, previous work has shown that PA1 can regulate proliferation to account for significant reductions in progenitor numbers [22]. Therefore, size of the neural crest progenitor population is not likely to explain the observed defects. Instead, the specificity of defects around the hinge region of PA1 suggest that reductions in *Fgf8* dosage disrupt jaw patterning.

In PA1, proximal-distal patterning is conferred by signals from the oral epithelia and *Fgf8* has previously been identified as a key contributor to this patterning [11, 14, 19, 23]. In particular, it has been hypothesized that patterning and polarity of the jaw are regulated by the interaction of signals emanating from the oral ectoderm and the first pharyngeal plate, which is formed where the ectodermal cleft contacts the first pharyngeal pouch endoderm [23, 24]. *Fgf8* is expressed in all three of these epithelia, and these tissues are underdeveloped and mis-aligned in mutant embryos (Fig. 6). The disruption of these tissue interactions is associated with alterations to pharyngeal morphology as well as the expression of key genes associated with pharyngeal patterning. Future work investigating how *Fgf8* regulates the morphogenesis of these epithelia will be important to understand how patterning of the jaw is established.

### 3.2 *Fgf8* dosage effects and phenotypic variation

We have previously described how *Fgf8* mediates non-linear dosage effects on craniofacial shape [7]. Here, we focus on the impact of *Ffg8* dosage on jaw development, and more specifically, mandibular development. We have found that reductions in *Fgf8* have a dosage dependent effect on jaw size, shape, and symmetry. On average, *Fgf8*^*Neo/Neo*^ E10.5 embryos express 35% of WT levels of *Fgf8*, while *Fgf8* ^*Δ/Neo*^ embryos express roughly 20% of WT Fgf8 levels [7]. This indicates that the entire range of morphological variation seen in the jaws of both mutants occurs within a molecular range 15% of *Fgf8* expression. On the other hand, the difference between *Fgf8* expression in WT and heterozygous (*Fgf8* ^*Δ/+*^) embryos is 40%, yet these two genotypes are indistinguishable morphologically. These data are consistent with the hypothesis that developmental defects exhibit non-linear genotype-phenotype relationships (Fig. 7E). Interestingly, some tissues appear to be more sensitive to reductions in *Fgf8* dosage.

We have shown that *Fgf8* ^*Neo/Neo*^ neonates have small but normal limbs, consistent with previous reports [9]. However, Fgf8 is required for limb development as tissue-specific loss of *Fgf8* in the AER of the developing limb results in reduction or loss of skeletal elements [25, 26]. Nonetheless, limb development is more robust to loss of *Fgf8* than is craniofacial development, likely due to compensation by other Fgfs expressed in the AER. Global *Fgf8* knock-outs fail to gastrulate, but 80% of *Fgf8* ^*Δ/Neo*^ embryos complete gastrulation [16], indicating that gastrulation has a low threshold requirement for *Fgf8*. In contrast, both heart and craniofacial development are more sensitive to *Fgf8* levels. Abu-Issa and colleagues (2002) describe abnormal heart development in 66% of *Fgf8* ^*Δ/Neo*^ mutants, while roughly one-third of these severe mutants have grossly normal hearts. Further, only 2% exhibit *situs inversus*, suggesting that these heart defects are related to heart-specific *Fgf8* expression and not associated with defective left-right axis formation deriving from gastrulation related *Fgf8* expression at the node [16]. In contrast, 100% of *Fgf8*^*Neo/Neo*^ (mild mutants) exhibit craniofacial defects including cleft palate [9, 16]. These data suggest that craniofacial development is particularly sensitive to *Fgf8* dosage (Fig. 7E). These differences in sensitivity may reflect different evolutionary pressures for canalization versus evolvability in tissue development. Selection favoring evolvability in craniofacial development may underlie the adaptive radiation of vertebrates, but may also result in increased susceptibility to disease [27].

### 3.3 Directional asymmetry and cardiopharyngeal developmental interactions

The relatively mild defects observed in some *Fgf8*^*Neo/Neo*^ mutants suggests that this genotype expresses a range of *Fgf8* that is near the upper part of the genotype-phenotype curve (teal box in Fig. 7E). Further, the directional asymmetry observed in this genotype is suggestive of subtle, but consistent, left-right differences in *Fgf8* expression (or Fgf signaling) in jaw development. We hypothesize that directional asymmetry in *Fgf8* mutants results from asymmetric contributions of *Fgf8* (or some other Fgf family member) to PA1 development from heart progenitor populations. The jaw and heart share progenitors within the pharyngeal mesoderm [28]. Thus, the directional asymmetry may derive from asymmetry in these migrating populations and/or subsequent heart morphogenesis. Heart looping results in the right side of PA1 experiencing more proximate relationships with the developing heart than the left side (Fig. 7A-D). Either of these tissue interactions may allow for asymmetric contributions of morphogens expressed in heart progenitors to PA1 development.

Similar directional trends occur in human craniofacial anomalies, and several craniofacial syndromes including Chromosome 22q11 deletion (22q11del) syndrome and Charge syndrome are characterized by both craniofacial and heart developmental defects [4–6]. Fgf signaling has been implicated in 22q11del syndrome [17, 29, 30], and may have more widespread involvement in facial dysmorphology and asymmetry. Further investigation of how signaling is shared between these populations may help elucidate both shared mechanisms in craniofacial and cardiac disease as well as asymmetries in congenital disorders.

### 3.4 *Fgf8* reduction models human syngnathia and TMJ defects

In humans, two broad categories of syngnathia exist: bony fusion of the maxillary and mandibular alveolar ridges and zygomatico-mandibular fusion [31–33]. In the majority of the latter cases, the TMJ is intact. We have observed a similar pattern in our *Fgf8* mutants. A role for *Fgf8* in syngnathia and jaw joint formation has previously been reported [34]. In that study, it was hypothesized that loss of *Foxc1* results in downregulation of *Fgf8* in the oral ectoderm to 80% of WT levels (although this may also have included *Fgf8* expression in the pharyngeal plate). Our data suggest that this is not a sufficient reduction to cause skeletal defects and the syngathia observed in *Foxc1* mutants is likely due to some other function of this gene. This would not be surprising given the difference in specific aspects of the fusion, which include hyperplasia rather than hypoplasia, and the broader expression of *Foxc1* in the proximal PA1 mesenchyme. Nonetheless, genetic interaction between *Fgf8* and *Foxc1*, and potentially many other genes expressed in the pharyngeal region, may further contribute to variation in syngnathia phenotypes. However, the particular model we present here, especially the mild mutant (*Fgf8*^*Neo/Neo*^), provides an excellent system for future investigations of molecular and cellular mechanisms underlying syngnathia and TMJ defects that closely model those observed in human disease.

## 4. Experimental Procedures

### 4.1 Experimental animals

The *Fgf8* allelic series involves three viable, adult genotypes that can be crossed to generate five different embryonic genotypes [9]. These mice and embryos were genotyped as previously reported [7, 9, 14]. Embryos were collected and staged based on the number of days after the observation of a postcoital plug at E0.5. Neonates (P0) were collected at birth. Mouse experiments were approved by the University of Massachusetts Lowell (Neo series) and the $University of Utah (CRE series) Institutional Animal Care and Use Committees. The tissue specific knock-out lines were generated and genotyped as previously described [21, 35]. Embryos were dissected on ice and fixed in 4% paraformaldehyde (PFA; for histology, gene expression and μCT) or 95% ethanol (for skeletal preparation).

### 4.2 Bone and cartilage staining

Differential bone and cartilage staining was performed using standard procedures [36]. Briefly, after ethanol fixation, embryos were skinned and then stained with 0.1% Alizarin Red and 0.3% Alcian Blue in ethanol for 3 days. After staining, the remaining non-skeletal tissue was digested with 1% KOH. Skeletal tissue was then cleared in glycerol prior to imaging.

### 4.3 Quantification of jaw and limb length

Dentary and limb bones were isolated from neonates after differential bone and cartilage staining. Isolated bones were imaged on a dissecting microscope at 1x magnification. The length of the bones was measured using arbitrary units. The average length of wild-type bones was set to 1 and then each individual bone was quantified relative to that average. Symmetry was determined by subtracting the length of the right bone from the left bone for each individual.

### 4.4 Micro-CT (μCT) imaging

For analysis of soft tissues, neonatal heads were stained using a 0.7% Phosphotungstic Acid (PTA) solution following published methods [37]. Imaging was performed on paraffin wrapped samples. For analysis of isolated bone, publicly available scans of neonatal heads were used from a previously published study [7]. Scans of PTA-stained embryos were visualized using Dragonfly software (version 2021.1 build 977, Object Research Systems). Scans of isolated bone were visualized with MeshLab.

### 4.5 Quantification of temporalis and coronoid volumes

Temporalis and coronoid volumes were quantified within the Dragonfly software. μCT scans of each head were read into the program as tiff stacks. The pitch and yaw were adjusted to achieve a coronal view for each sample. A region of interest (ROI) mask was created separately for each measured structure of each sampled head. The round brush within the ROI Painter tool was used to label each voxel corresponding to the sections of each separate structure(s). By choosing separate masks and labeling the structure’s voxels, a 3D image can be created for each structure, based on the morphology within the sample. Once the voxels of each structure were labeled, the Dragonfly software automatically calculates the total number of voxels labeled (based on the ROI mask chosen) and gives a volume of the chosen mask. By using the underlying morphologies to dictate which voxels were labeled, we used this method to calculate 3D volumes of the temporalis and coronoid within each sample given as mm^3^. One-way ANOVAs were used to evaluate the difference between genotypes and side.

### 4.6 Histology/Trichrome Staining

Neonatal and E14.5 heads were dehydrated in ethanol after overnight PFA fixation. The heads then were processed with Clear-Rite, a xylene substitute, to remove ethanol, and finally embedded in paraffin (Surgipath Paraplast Plus, Leica) inside a vacuum drying oven (ADP 31, Yamato). Embryo heads were sectioned at 10um using a microtome. Sections were stained using Milligan’s trichrome [38], mounted with Permount, and imaged under a bright field microscope.

### 4.7 *in situ* hybridization

Probes for *in situ* hybridization were generated from RNA isolated from E9.5 and E10.5 embryos. cDNA was produced using the Maxima first strand synthesis kit (ThermoFisher, K1641). The cDNA was then used as a PCR template to amplify the gene of interest (GOI). Select PCR primers had linkers (Fw:5’-ggccgcgg-3’; Rv:5’-cccggggc-3’) to allow for a nested PCR TOPO cloning. PCR purification of these templates used a gene specific forward primer (GSFP) and the T7 Universal primer to amplify the initial GOI template. Primers lacking linkers were TOPO cloned into TOP10 competent E. coli cells using a PCR4 topo vector (ThermoFisher). Colonies were screened to ensure correct band length. Once purified, samples were sent to Genewiz to be sequenced to ensure the correct GOI was amplified and direction of the insertion. A second PCR was used to amplify the GOI using M13 primers after plasmid verification. Second PCR product lengths were verified on a 1% agarose gel. Products from the secondary PCR were used as the template for the anti-sense mRNA probe. Probes were made by using a dig RNA labeling mix (Sigma-Aldrich, #11277073910). Fluorescent probes were created with A555 fluorescent tyramine following publicly available protocols (https://sites.google.com/site/helobdellaprotocols/histology/tsa). Fluorescent samples were counter-stained using Hoechst’s O.N. (1:1000 dilution of 10 mg/ml stock) and were imaged using a 40x oil immersion objective lens on a Leica sp8 confocal microscope.

### 4.8 Apoptosis staining

Apoptotic cells were identified using Lysotracker Red (ThermoFisher). Unfixed E9.5 or E10.5 embryos were stained for 30 minutes in Lysotracker solution as previously described [39]. Stained embryos were clarified with methanol and then fixed in 4% PFA.

## 5 Acknowledgements

We would like to thank Evelyn Schwager for technical support. This work was supported by the National Institutes of Health: R03 DE028984 to JLF and R01 DE019638 to BH and RSM.

## Figure Legends

**SupFig1:**
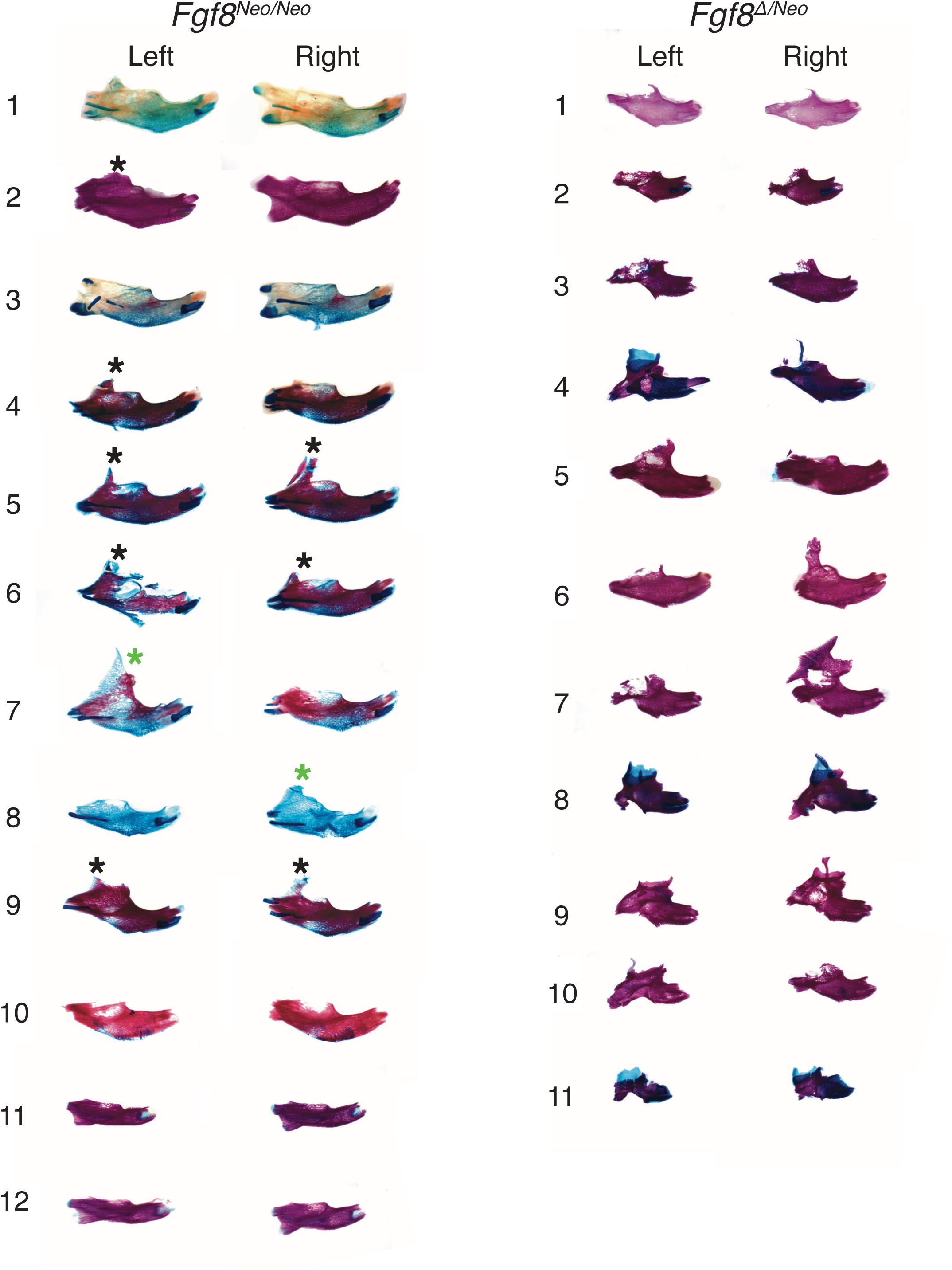
Fgf8 mutant mandibles exhibit inter- and intra-individual variation in severity. Isolated dentary bones differentially stained for bone (red) and cartilage (blue) from *Fgf8*^*Neo/Neo*^ (left) and *Fgf8*^*Δ/Neo*^ (right) neonates. We observe unilateral fusion in 11/34 *Fgf8*^*Neo/Neo*^ jaws (including data from microCTs). Black asterisks indicate mild fusion, with jugal-zygomatic process present; green asterisks indicate more severe fusion.

**SupFig2:**
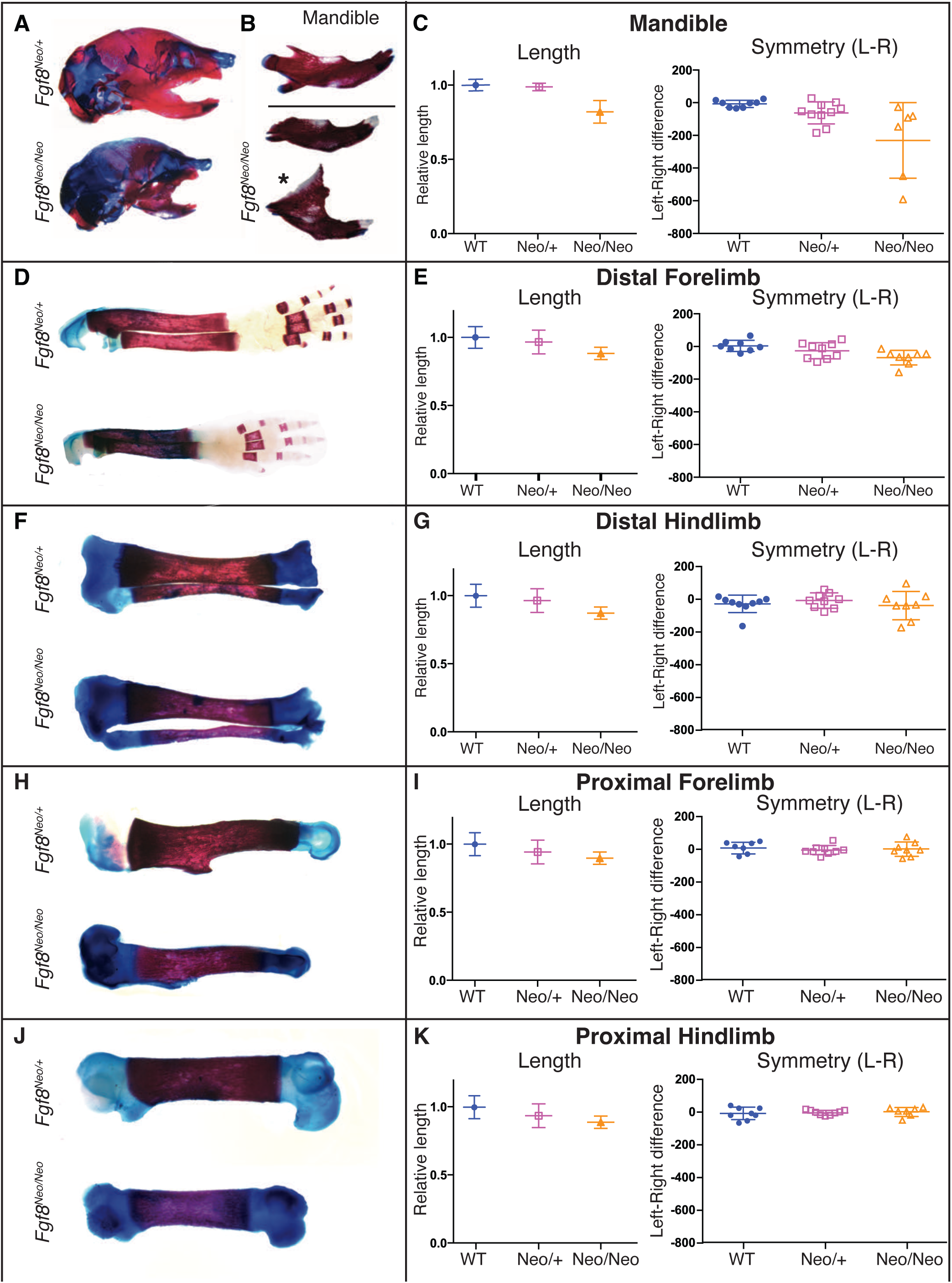
*Fgf8*^*Neo/Neo*^ mandibles exhibit directional asymmetry. **A)** Neonatal (P0) skulls differentially stained for bone (red) and cartilage (blue) from *Fgf8*^*Neo/+*^ (upper) and *Fgf8*^*Neo/Neo*^ (lower) neonatal mice show hypoplasia of the jaw in mutant embryos. **B)** Isolated dentary bones differentially stained for bone (red) and cartilage (blue) from *Fgf8*^*Neo/+*^ (upper) and *Fgf8*^*Neo/Neo*^ (lower) neonatal mice. **C)** The length (left) of isolated dentaries were measured using arbitrary units and symmetry (right) was determined by subtracting the length of the right side from the length of the left side for each individual. **D-K)** Isolated limb bones differentially stained for bone (red) and cartilage (blue) from *Fgf8*^*Neo/+*^ (upper) and *Fgf8*^*Neo/Neo*^ (lower) neonatal mice were measured for length (left) and symmetry (right). n= 9 per genotype.

**SupFig3:**
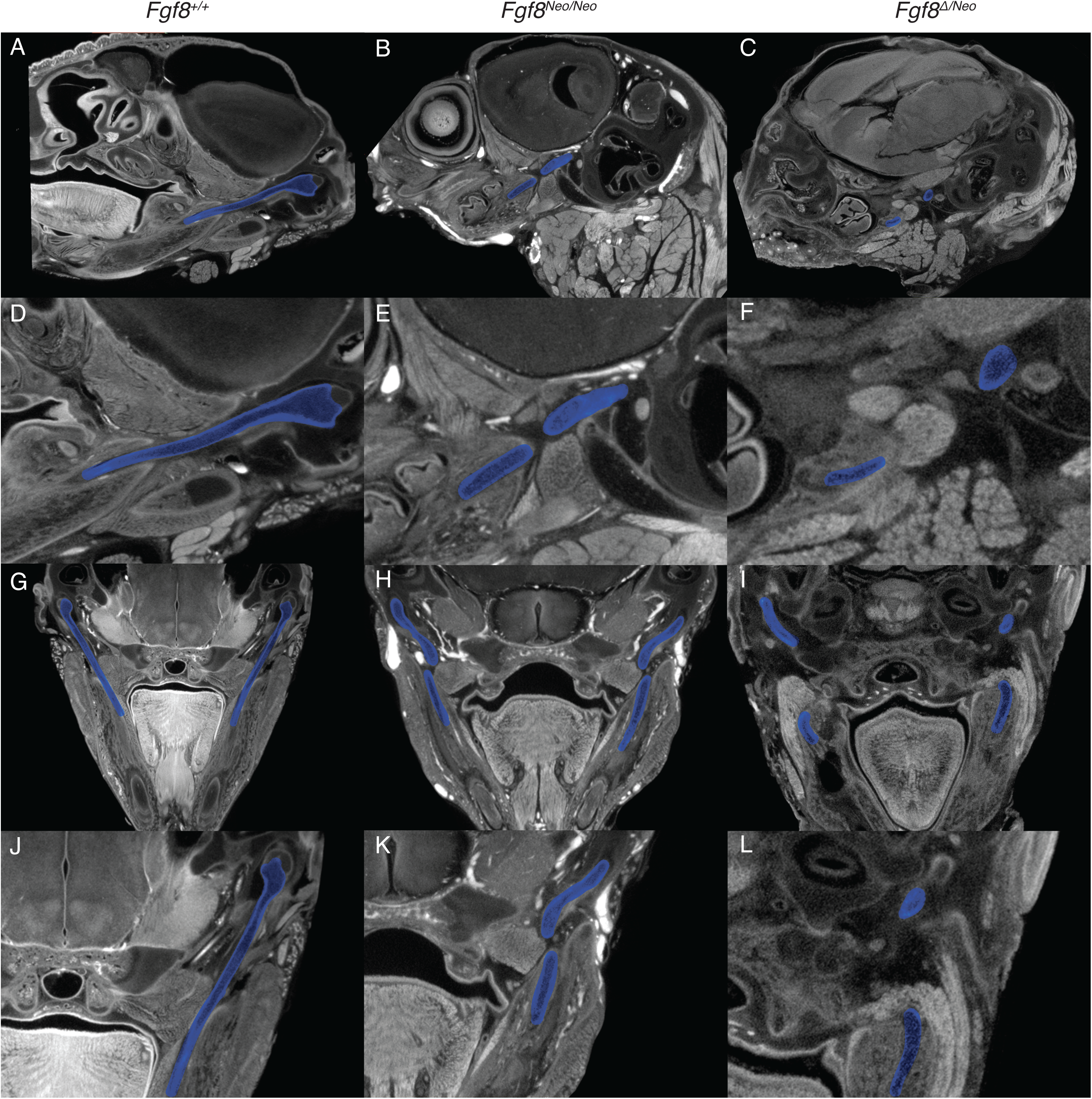
*Fgf8* mutants exhibit discontinuity of Meckel’s cartilage. Mirco-CT scans of neonatal A,G) *Fgf8*^*+/+*^ (WT), B,H) *Fgf8*^*Neo/Neo*^, and C,I) *Fgf8*^*Δ/Neo*^ skulls demonstrate that Meckel’s cartilage (MC; blue) is discontinuous in mutants. D-F, J-L) Enlarged images illustrate the left and right sides of MC. The distance between the distal and proximal segments of MC is greater in *Fgf8*^*Δ/Neo*^ than in *Fgf8*^*Neo/Neo*^ suggesting a dosage effect. In both mutants, MC is present in a proximal (middle ear) and distal (dentary) segment. The break in MC occurs proximal to where the Inferior Alveolar Nerve branches.

**SupFig4:**
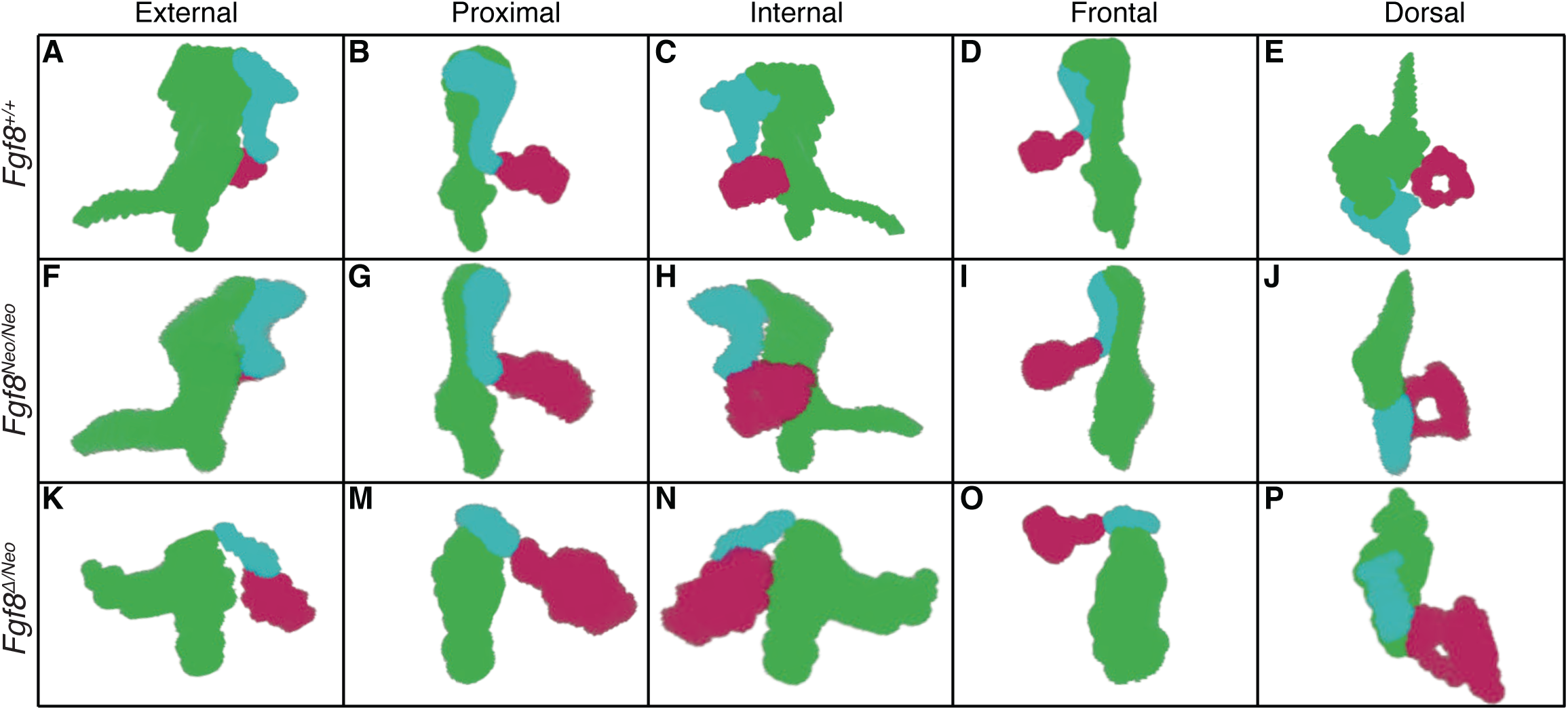
*Fgf8* dosage levels influence malleus size, rotation, and manubrium angle. Three-dimensional reconstructions of the malleus (green), incus (blue) and stapes (pink) of A-E) *Fgf8*^*+/+*^ (WT), F-J) *Fgf8*^*Neo/Neo*^ and K-P) *Fgf8*^*Δ/Neo*^ neonatal skulls. F-I) The manubrium of the malleus (mM) in *Fgf8*^*Neo/Neo*^ is more perpendicular relative to the process brevis (mPB) and curves upwards in *Fgf8*^*Δ/Neo*^ mice. In the first 4 columns, mPB is used as a vertical reference, therefore, the malleus in *Fgf8*^*Neo/Neo*^ is rotated shifting the head of the malleus rostrally, which is further evident in *Fgf8*^*Δ/Neo*^.

**SupFig5:**
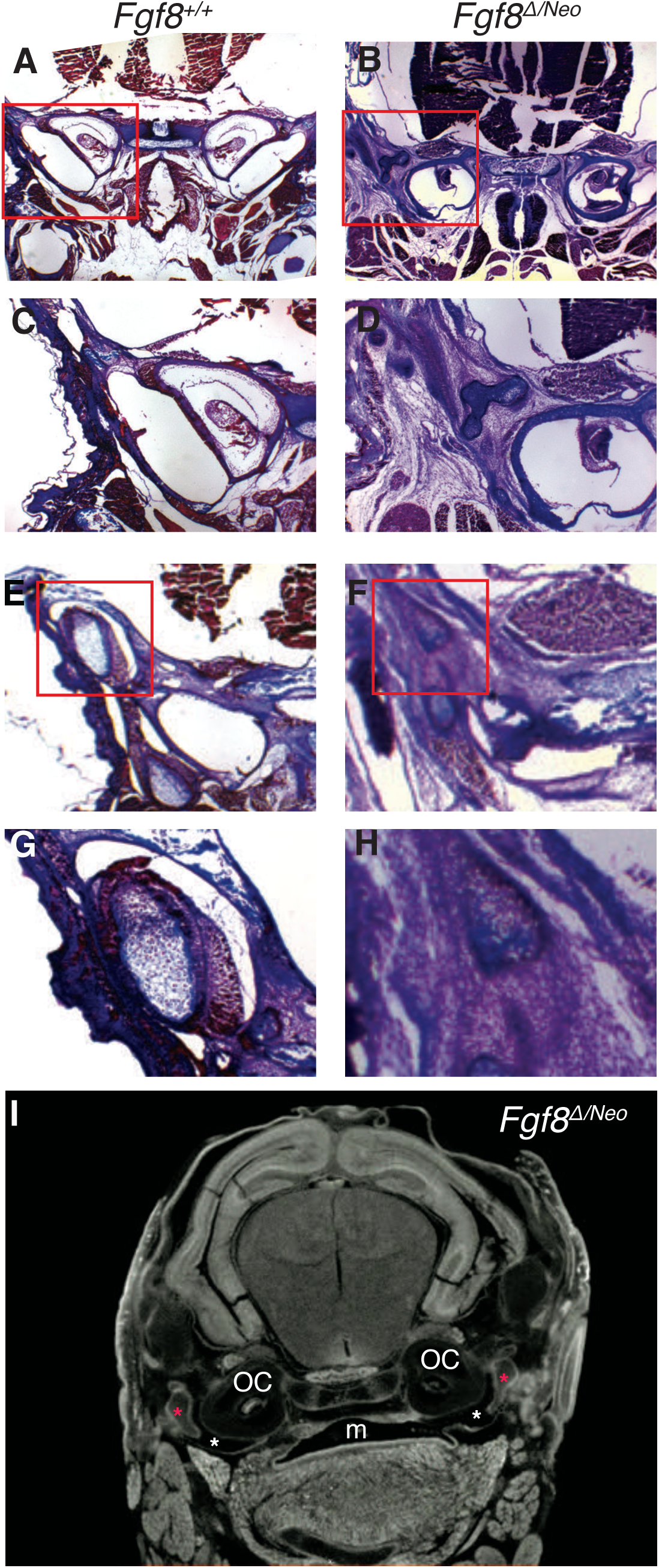
Severe reductions of *Fgf8* deform the temporomandibular jaw joint (TMJ). Coronal trichrome stained cross-sections of A) *Fgf8*^*+/+*^ (WT) and B) *Fgf8*^*Δ/Neo*^ neonatal (P0) skulls. Panels A-D are more proximal in the head than panels E-H. A) WT neonatal mice have large tympanic cavities separating the malleus and cochlea. B) *Fgf8*^*Δ/Neo*^ have variable defects of the middle ear. In this example, the tympanic cavity is absent on the left side, but only reduced on the right. Red boxes in A and B are shown magnified in panels C and D. E) In WT, the TMJ has a joint disc below a defined glenoid fossa creating an upper joint cavity depicted in panel G. F) *Fgf8*^*Δ/Neo*^ neonates have a severely reduced the glenoid fossa (GF) and condylar process, the joint cavity is malformed, and the articular disc is absent shown in panel H. I) micro-CT scans of *Fgf8*^*Δ/Neo*^ also show that the mouth cavity can open into the tympanic cavities of the ear (white asterisk) but the presence of the malleus (red asterisks) is present on both sides. disk (D), mouth (M), otic capsule (OC), upper joint cavity (UJV). (total stained embryos: *Fgf8*^*+/+*^ (WT) 4; *Fgf8*^*Δ/Neo*^ 3)

**SupFig6:**
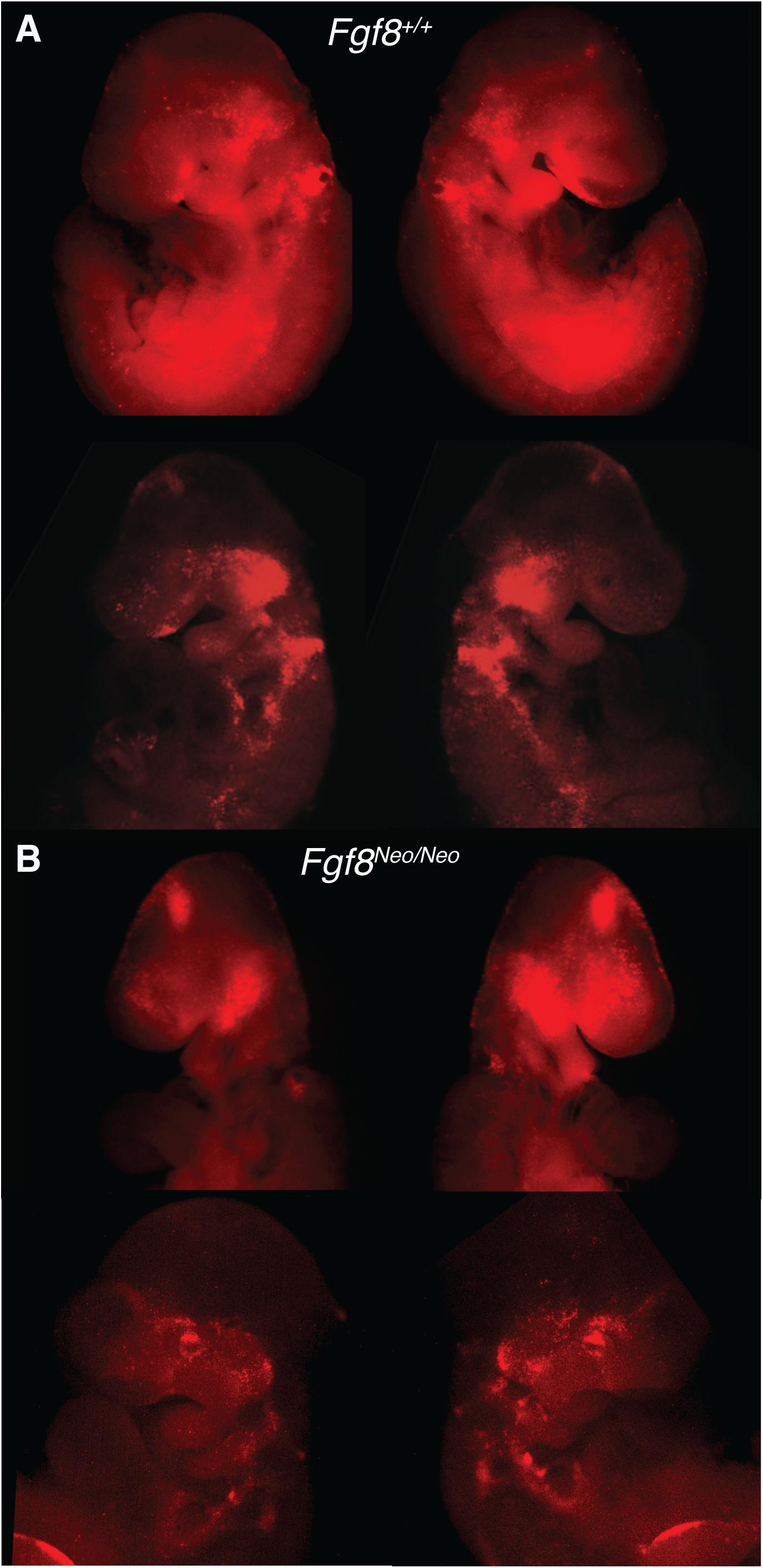
Cell death is altered, but not asymmetric *Fgf8*^*Neo/Neo*^ embryos. Lysotracker (red) staining of apoptotic cells is shown in 2 separate E9.0 A) WT and B) *Fgf8*^*Neo/Neo*^ embryos. Representative images are provided for each genotype (total stained embryos: *Fgf8*^*+/+*^ (WT) 12, *Fgf8*^*Neo/+*^ 11, *Fgf8*^*Neo/Neo*^ 7).

**SupFig7:**
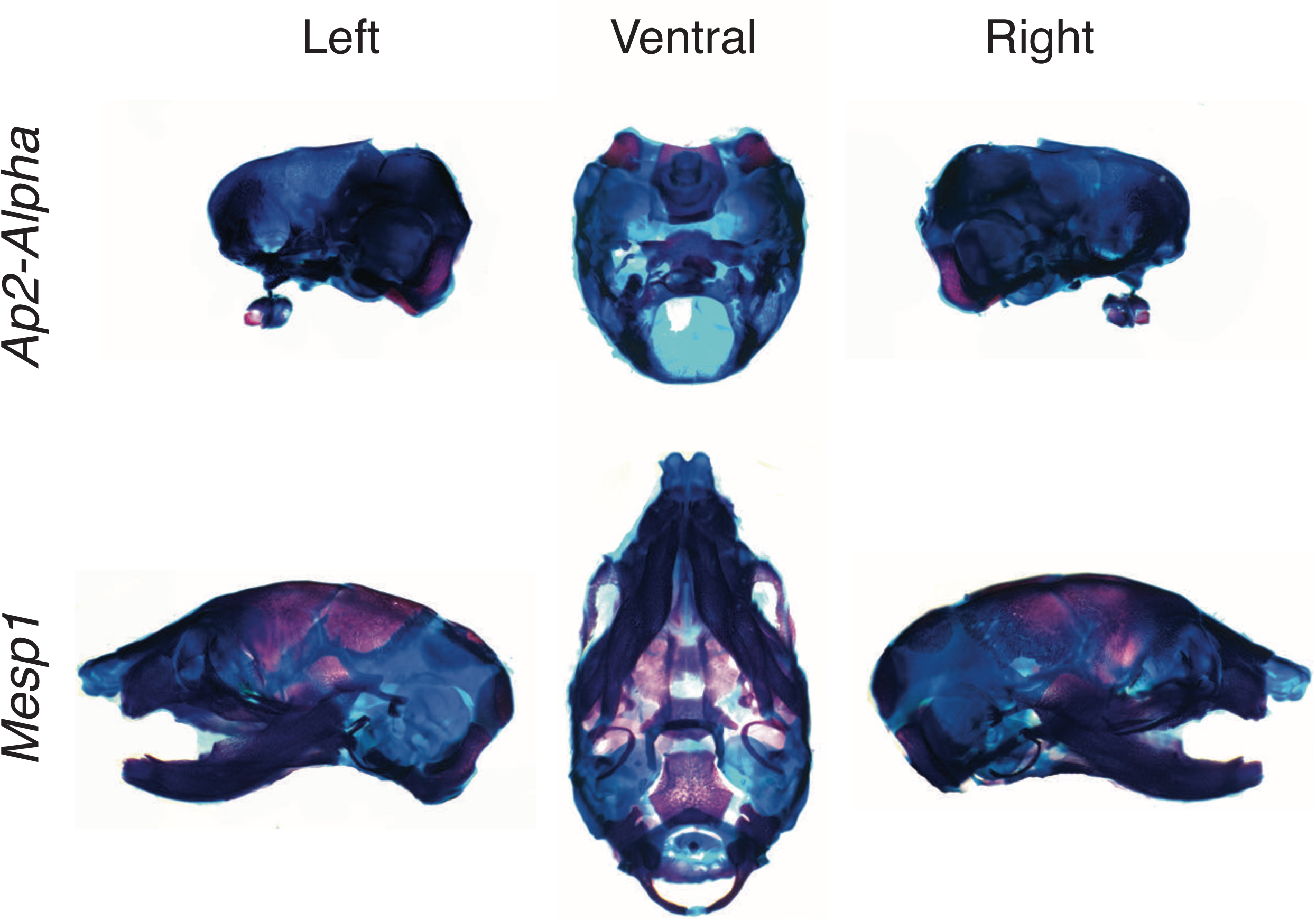
Ectodermal and mesodermal specific loss of Fgf8 do not alter asymmetry patterns. Neonatal skulls differentially stained for bone (red) and cartilage (blue). Representative images are shown for tissue-specific Fgf8 knock-outs using *Ap2-alpha* CRE (upper) and *Mesp1* CRE (lower). Representative images are shown for 3 *Ap2-alpha* mutants and 5 *Mesp1* mutants.

